# The role of world knowledge in sentence comprehension under working-memory constraints: An fMRI study

**DOI:** 10.64898/2026.10.06.756870

**Authors:** Rei Emura, Hyeonjeong Jeong, Yousuke Kawachi, Ryuya Komuro, Masatoshi Koizumi

## Abstract

**Introduction:** Comprehension of thematic roles, i.e., “who did what to whom,” relies on limited cognitive resources, and constraints on these resources can alter the use of plausibility information derived from world knowledge. Although sentence-processing studies have described how plausibility information constrains interpretation, it remains unclear how this process is supported by the architecture of working memory. This study examines whether the phonological loop, one of the components of working memory, contributes to thematic-role processing based on world knowledge.

**Methods:** Using articulatory suppression and functional magnetic resonance imaging, we examined whether constraining the function of the phonological loop modulates behavioral and neural responses to sentences in which syntactically assigned thematic roles are plausible or implausible given world knowledge. To constrain the phonological loop, we manipulated the task: a single task and a dual task involving concurrent articulation.

**Results:** We found no significant interaction between task and plausibility, although equivalence analyses did not provide evidence for the absence of a plausibility effect during articulation. In contrast, a robust main effect of thematic-role plausibility was observed: implausible sentences elicited greater activation than plausible sentences in the left inferior frontal gyrus (LIFG) and medial superior frontal gyrus (mSFG) at the whole-brain level, and in both the pars triangularis and pars orbitalis of the LIFG in the ROI analyses.

**Discussion:** These findings provide no conclusive evidence that constraining the phonological loop substantially modulates the use of world knowledge in comprehension of thematic roles, while also leaving open the possibility of a smaller task-dependent effect. At the same time, the robust neural responses to implausibility indicate that the LIFG and mSFG consistently contribute to the processing of plausibility information derived from world knowledge.

## 1. Introduction

Comprehension of thematic roles, i.e., “who did what to whom,” depends not only on syntactic structure but also on plausibility derived from world knowledge (Bates et al., 1982; Gibson and Pearlmutter, 1998; MacWhinney and Bates, 1989). For example, consider the following sentences:

1. Plausible: *The baby was held up by her mother*.
2. Implausible: *The mother was held up by her baby*.

In an implausible sentence such as (2), the syntactically correct parse, in which *the baby* is the agent and *her mother* is the patient, conflicts with the more plausible interpretation that the mother held up the baby. This conflict gives rise to competition between syntax-based and plausibility-based processes (MacWhinney and Bates, 1989), as evidenced by longer reading times (Rayner et al., 2004), a higher rate of misinterpretations (Ferreira, 2003), a posterior positivity in event-related potentials (Ye et al., 2007), and increased brain activity in regions such as the left inferior frontal gyrus (LIFG) (Hagoort et al., 2004; Sassa et al., 2007; Ye and Zhou, 2009).

Importantly, the processing of plausibility information is supported by working memory, as evidenced by the finding that people rely more on plausibility than on syntax when cognitive resources are limited. For example, the difference in accuracy between implausible and plausible sentences is smaller for syntactically complex sentences than for simple sentences (Andrews et al., 2017; Ayasse et al., 2021; Ferreira, 2003; Gibson et al., 2013). Increased reliance on plausibility has also been observed in populations with limited memory resources, including people with lower verbal working memory capacity (Andrews et al., 2017), people with aphasia (Gibson et al., 2016), second-language learners (Futrell and Gibson, 2017), and older adults (Christianson et al., 2006; Liu, 2024). These findings suggest that the relative contributions of syntactic and plausibility information varies with the availability of working memory resources.

These behaviors have been accounted for by the rational inference approach, or noisy-channel theory (Gibson et al., 2013, 2016). According to this account, limited cognitive resources may increase noise in the internal representation of linguistic input, thereby increasing uncertainty about the reliability of syntactic evidence. As a result, comprehenders place relatively greater weight on prior knowledge, such as semantic plausibility, than on the syntactic cues in the input.

However, it remains unclear which specific component of working memory contributes to integrating plausibility information into comprehension of thematic roles. Sentence comprehension relies on finite-capacity working memory resources for both storage and processing (Baddeley, 2003, 1979; Just and Carpenter, 1992). The classical working memory model, known as the “three-component model,” divides working memory into three subsystems: the central executive, an attentional control system with limited capacity; the visuo-spatial sketchpad, which handles visual and spatial information; and the phonological loop (Baddeley, 1979). The revised model, called “multi-component model,” adds the episodic buffer as a multimodal store specialized for holding integrated representations (Baddeley, 2000; Hitch et al., 2020, for reviews). The phonological loop specifically supports incremental sentence processing by maintaining verbal input through subvocal articulation, thereby preventing its decay. During sentence comprehension, world knowledge is retrieved from long-term memory and used together with information derived from the linguistic input. According to the theories, the phonological loop may support the processing of plausibility information, but it has not been examined yet.

Taken together, whereas the rational-inference account characterizes sentence interpretation in terms of the contributions of linguistic information and prior knowledge, working-memory accounts provide a framework for asking how information required for this interpretation may be maintained within working memory. Thus, although plausibility (in)consistency is known to be sensitive to resource limitations, it is still unclear whether these effects arise from phonological-loop constraints or from other aspects of working memory. If the phonological loop supports the maintenance of incoming verbal input during the integration of world knowledge and linguistic input, then disrupting the phonological loop should affect how plausibility information is used during the comprehension of thematic roles.

One way to test this hypothesis is to examine whether neural responses associated with sentence processing are sensitive to constraints on the phonological loop. A functional magnetic resonance imaging (fMRI) study of syntactic processing has distinguished neural responses that remain specific to linguistic processing from those that vary with broader working-memory demands (Rogalsky et al., 2008). In particular, the study compared the processing of syntactically simple subject-relative sentences and complex object-relative sentences under single-task and concurrent articulation conditions (referred to as “articulatory suppression”). In the single-task condition, complexity increased activation in both LIFG pars opercularis (LIFGoper) and LIFG pars triangularis (LIFGtri). However, when participants concurrently articulated “da ba ga da…” while listening to sentences, the complexity effect remained in LIFGtri but was reduced in LIFGoper. This dissociation has been interpreted as evidence that LIFGtri is more specifically involved in syntactic processing, including the retrieval and manipulation of syntactic structures, whereas LIFGoper is more sensitive to phonological-loop demands (Matchin, 2018). More broadly, neuroimaging studies have distinguished language-specific and domain-general responses within the LIFG (Bulut, 2023; Fedorenko et al., 2011, 2012; Fedorenko and Blank, 2020; Nozari and Martin, 2024). These findings suggest that manipulating phonological-loop resources while measuring brain activity can help determine whether a sentence-processing response is maintained independently of phonological demands or is modulated by them. Following this approach, we use this articulatory-suppression paradigm to examine the neural consequences of constraining the phonological loop during sentence comprehension based on world knowledge.

A similar approach may be informative for thematic-role interpretation based on world knowledge. The LIFG is associated with both syntactic processing and working-memory demands (Caldwell, 2026; Fedorenko et al., 2011; Matchin, 2018; Matchin et al., 2024; Rogalsky and Hickok, 2011), and its subregions have been argued to differ in their sensitivity to linguistic and domain-general demands (Bulut, 2023; Fedorenko et al., 2011, 2012; Fedorenko and Blank, 2020; Nozari and Martin, 2024). The LIFG has also been implicated in both phonological-loop processing (Emch et al., 2019; Nishida et al., 2017) and the processing of sentences in which syntactically specified interpretations are inconsistent with world knowledge (Hagoort et al., 2004; Kuperberg et al., 2008; Sassa et al., 2007; Tyler et al., 2010; van de Meerendonk et al., 2013; Ye and Zhou, 2009). In particular, these studies have consistently reported involvement of LIFGtri and LIFG pars orbitalis (LIFGorb). However, unlike the syntactic-processing study by Rogalsky et al. (2008), these studies did not directly constrain phonological-loop resources. We therefore focus on LIFGtri and LIFGorb and examine whether neural responses to thematic-role plausibility vary under constraints on the phonological loop.

To summarize, the objective of this study is to examine whether the integration of world knowledge into thematic-role comprehension is supported by the phonological loop, one of the components of working memory. Specifically, this study examines whether phonological-loop constraints modulate neural responses to thematic-role plausibility. In the fMRI experiment, participants listened to plausible and implausible sentences under a single-task condition, in which they focused solely on sentence comprehension, and an articulatory-suppression condition, in which they concurrently articulated nonwords while listening to the sentences (see Table 1). We expected lower comprehension accuracy and greater activation in LIFGtri and LIFGorb for implausible than for plausible sentences, reflecting the thematic-role plausibility effect. The critical question was whether this plausibility effect would differ between the single-task and articulatory-suppression conditions. If the phonological loop contributes to the processing of plausibility information, a reduced plausibility effect under articulatory suppression should be observed.

**Table 1:**
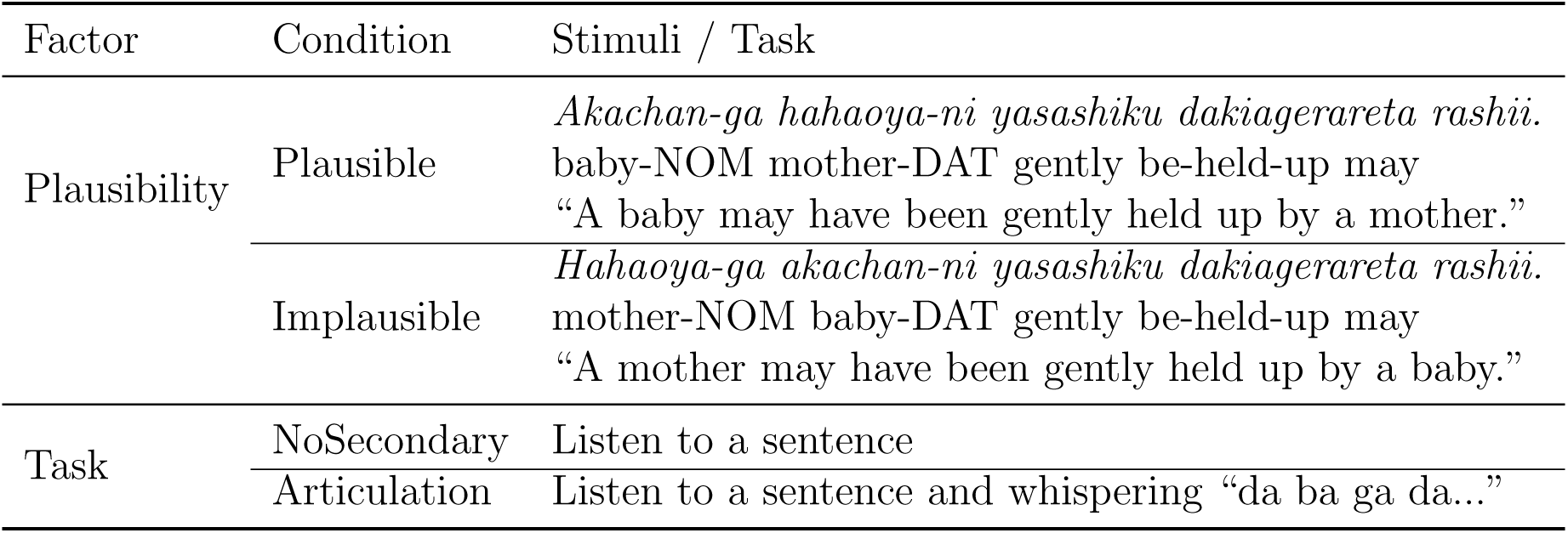
Experimental design. Plausibility shows stimuli, and the secondary task shows the task procedure. NOM = Nominative case; DAT = Dative case.

| Factor | Condition | Stimuli / Task |
| --- | --- | --- |
| Plausibility | Plausible | *Akachan-ga hahaoya-ni yasashiku dakiagerareta rashii.*<br>baby-NOM mother-DAT gently be-held-up may<br>“A baby may have been gently held up by a mother.” |
|  | Implausible | *Hahaoya-ga akachan-ni yasashiku dakiagerareta rashii.*<br>mother-NOM baby-DAT gently be-held-up may<br>“A mother may have been gently held up by a baby.” |
| Task | NoSecondary | Listen to a sentence |
|  | Articulation | Listen to a sentence and whispering “da ba ga da...” |

## 2. Materials and Methods

### 2.1. Participants

This study included 62 right-handed native Japanese speakers. None of the participants reported any history of neurological or psychiatric problems. One participant was excluded due to low accuracy on comprehension questions (less than 80% accuracy on plausible and filler stimuli), one participant due to excessive head movement (greater than 5 mm), and two participants due to excessive head movement and rotation (greater than 10*^◦^*). Thus, the final sample included 58 participants (20 females and 38 males; mean *±* SD age: 21.69 *±* 3.32 years). Based on a Japanese translation of the Edinburgh Handedness Inventory with ten items (Oldfield, 1971), all the participants were right-handed (mean laterality quotient: 97.30 *±* 5.62). We obtained written informed consent from all participants before participation in the experiment, and this study was approved by the Ethics Committee of the Smart-Aging Research Center and the Graduate School of Arts and Letters, Tohoku University. This study was conducted in accordance with the guidelines of the Declaration of Helsinki. The participants received 3,000 JPY for their participation.

### 2.2. Design and stimuli

We used a 2 × 2 factorial design manipulating the secondary task (referred to as “task”) and the plausibility of the stimuli (see Table 1). Tasks included the NoSecondary condition (participants only listened to sentences) and the Articulation condition (participants listened to sentences while whispering “da ba ga da…”). The plausibility factor comprised the Plausible condition and the Implausible condition, exemplified in Table 1. All target sentences were in the passive voice, and implausible stimuli were generated by swapping the subjects and objects in the plausible stimuli. Stimuli were selected based on a norming study, and their plausibility was assessed in a follow-up questionnaire (see the Supplementary Material).

In total, we created 24 stimuli for each plausibility. We constructed two lists to avoid presenting identical stimuli across task conditions within a list. Condition assignments were counterbalanced. Each list contains 48 target stimuli and 48 filler stimuli. In this experiment, we included plausible sentences with a variety of syntactic structures as filler stimuli, and intermixed them with the target stimuli. Each list was presented across four sessions. The order of stimuli was randomized within each session.

We matched the length of the target sentences as much as possible. The mean audio duration was 3.95 seconds (from 3.43 to 4.31 seconds), and the mean number of morae was 23.88 (from 20 to 26 morae) in the target stimuli. Filler stimuli had a mean duration of 3.80 seconds (from 3.15 to 4.27 seconds) and a mean of 22.31 morae (from 14 to 26 morae).

### 2.3. Procedure

We conducted the sentence comprehension task as follows (illustrated in Figure 1). First, an instruction was presented on the display; participants immediately began whispering “da ba ga da…” if the articulation condition was assigned. We instructed them to say “daba” and “gada” in time with a metronome set to a tempo of 145 beats per minute (the metronome was not used during the experiment). After 1,000 ms, participants heard a sentence. When the sentence ended, a fixation cross appeared for a jittered interval of 2,000–4,000 ms, and participants stopped whispering in the articulation condition. Finally, they answered a two-choice question related to the sentence. The questions were about who did the action in the target stimuli, and about broad questions (e.g., who, to whom, where, and how) in the fillers. The place of the answer was counterbalanced. Feedback was not given.

**Figure 1:**
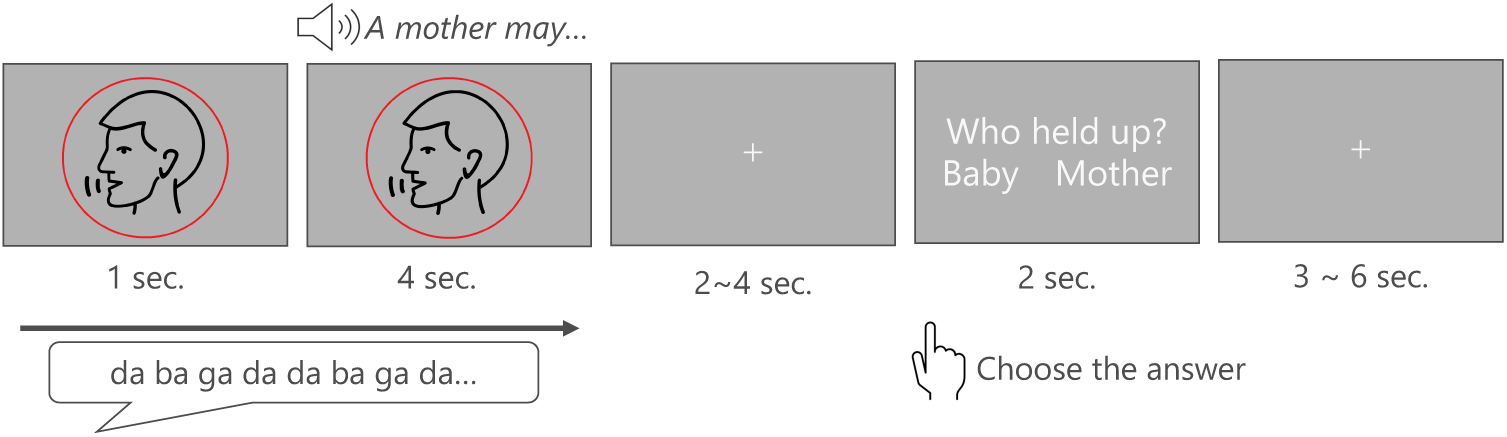
Procedure for the experiment. In the Articulation condition, participants were asked to whisper “da ba ga da…” while listening to a sentence. In the NoSecondary condition, they just listened to a sentence. In both conditions, they answered a two-choice question after each sentence. Stimuli and questions were provided in Japanese.

After the fMRI experiment, we administered the operation span task (Christianson et al., 2006) to measure participants’ working memory capacity (WMC), using a Japanese version of PySpanTask (von der Malsburg, 2015). The procedure of the operation span task is as follows. At first, participants read aloud each presented equation and verified its correctness. After a response was made or the time limit was reached, a single alphabet letter was presented for 800 ms. Participants were asked to remember the letter. After 3–8 equation–letter pairs, participants were asked to type the letters in the order of presentation. There were 32 trials, and the presentation order was randomized. Before the actual test, participants practiced with four trials. Partial-credit unit scores, which are the mean proportion of correctly recalled items, were used as the WMC score as recommended by Christianson et al. (2006).

### 2.4. Imaging acquisition and preprocessing

fMRI images were acquired using a Philips Achieva dStream 3.0T MRI scanner (Eindhoven, The Netherlands). Task stimuli were presented via PsychoPy (Peirce et al., 2019). Functional images were collected using an echo-planar imaging (EPI) with the following parameters: repetition time (TR) = 1,600 ms; echo time (TE) = 30 ms; flip angle (FA) = 80°; field of view (FOV) = 192 mm; voxel size = 3.00 *×* 3.05 *×* 4.00 mm^3^; slice thickness = 4 mm; slice gap = 0 mm. Twenty-six axial slices covering the entire brain were obtained at 1.6-s intervals. In total, 297 volumes were acquired from each participant during each of the four sessions. Subsequently, T1-weighted magnetization-prepared rapid gradient-echo (MPRAGE) images were also acquired with the following parameters: TR = 11 ms; TE = 5.1 ms; FA = 8°; FOV = 256 mm; voxel size = 0.70 mm.

Functional images were preprocessed using Statistical Parametric Mapping software (SPM12; Wellcome Centre for Human Neuroimaging, London, UK) running in the MATLAB environment. Preprocessing included slice-timing correction, realignment for head-motion correction, co-registration of each participant’s T1-weighted structural image to the mean functional image, normalization to Montreal Neurological Institute (MNI) space, and spatial smoothing with an 8-mm full-width at half-maximum (FWHM) Gaussian kernel.

### 2.5. Data analysis: behavior

For the behavioral data, we evaluated accuracy for the comprehension task in each condition, and conducted a statistical test using generalized linear mixed-effects (GLM) models using the lmerTest package (Kuznetsova et al., 2017) in R (R Core Team, 2025).^1^ The model included secondary task and plausibility as fixed effects and random intercepts for participants and items.

### 2.6. Data analysis: brain activity

For the brain data, we employed a conventional two-level approach for event-related fMRI analysis using SPM12. In the first-level analysis within the general linear framework, we conducted voxel-wise multiple regression analyses. This analysis was based on the expected signal changes for each condition constructed using SPM12’s hemodynamic response function. These models were applied to the preprocessed images of each participant. The activation levels were estimated using a multi-session design matrix that modeled four conditions: NoSecondary–Plausible, NoSecondary–Implausible, Articulation–Plausible, and Articulation–Implausible.

Each trial in every condition was modeled at the onset of the verb of the target sentence with a duration of the verb (end of sentence) plus 2,000 ms because participants are expected to detect the plausibility of sentences at the timing of the verb. We also modeled the onset of the beginning of the sentence with a duration of the whole sentence plus 2,000 ms. Contrast images were created for each participant under each condition and between conditions.

To reduce motion-derived artifacts, we included six motion parameters (three translations and three rotations) as nuisance regressors. In addition, we applied motion censoring, by including a volume-wise spike regressor that was coded 1 for volumes exceeding the framewise displacement (FD) threshold of 0.5 and 0 otherwise (Power et al., 2012; Siegel et al., 2014). We then calculated FD values for each condition from the six realignment parameters, with rotational displacements converted to millimeters assuming a 50-mm head radius. Mean FD values were 0.20 *±* 0.10 (mean *±* standard deviation) for NoSecondary–Plausible, 0.20 *±* 0.10 for NoSecondary–Implausible, 0.34 *±* 0.15 for Articulation–Plausible, and 0.34 *±* 0.15 for Articulation–Implausible. According to a linear mixed-effects (LME) model including task, plausibility, and their interaction as predictors, with participant included as a random intercept, there was no significant effect of plausibility (*β* = 0.00, SE = 0.00, *t* = 0.64, *p* = 0.520), although there was a significant effect of task (*β* = *−*0.11, SE = 0.00, *t* = *−*27.11, *p <* 0.001). The interaction was not significant (*β* = 0.00, SE = 0.00, *t* = *−*0.02, *p* = 0.986). In sum, motion-related confounds were unlikely to differ systematically across conditions, particularly the plausibility effect.

The second-level analysis involved statistical inference on contrasts in the parameter estimates. This was achieved using a between-participants (random effects) model evaluated using one-sample t-tests. First, to examine the brain activation induced by the type of secondary task, Articulation conditions were compared with NoSecondary conditions. Second, to investigate the brain areas associated with plausibility, Plausible conditions were compared with Implausible conditions. Third, to examine the interaction effect between the type of secondary task and the type of plausibility, interaction contrasts were defined as [(Articulation: Implausible *>* Plausible) *>* (NoSecondary: Implausible *>* Plausible)] and [(NoSecondary: Implausible *>* Plausible) *>* (Articulation: Implausible *>* Plausible)], which were tested separately. Finally, to examine the plausibility effect within each secondary-task condition, simple-effect contrasts of Implausible *>* Plausible were tested separately for the NoSecondary and Articulation conditions.

The statistical threshold in the voxel-by-voxel analysis, assuming a search area of the whole brain, was set at uncorrected *p <* 0.001 (voxel level) and then cluster-level family-wise error (FWE) corrected to *p <* 0.05 (cluster size). For the post-hoc analyses examining the simple effect of plausibility within each task condition, Bonferroni correction was applied for the two comparisons. Accordingly, the threshold was set at voxel-level uncorrected *p <* 0.001, followed by cluster-level FWE correction at *p <* 0.025. Activation peak coordinates were reported in MNI space, and activated brain regions were identified using the automated anatomical labeling atlas (Tzourio-Mazoyer et al., 2002).

Moreover, we conducted literature-based region-of-interest (ROI) analyses using the MarsBaR toolbox (Brett et al., 2002) implemented in SPM12. We selected two ROIs in the left IFG: LIFGtri and LIFGorb, where previous work consistently shows greater activation of implausible sentences than of plausible sentences (Hagoort et al., 2004; Kuperberg et al., 2008; Sassa et al., 2007; Tyler et al., 2010; van de Meerendonk et al., 2013; Ye and Zhou, 2009). Spherical ROIs (8-mm radius) were defined around the peak voxels [-52, 26, 20] and [-48, 34, −2] in LIFGtri and LIFGorb, respectively, which are reported in a previous Japanese fMRI study (Sassa et al., 2007).

For each ROI, activation estimates were analyzed using a LME model with plausibility, task, and their interaction as fixed effects and participant as a random intercept by using the lmerTest package (Kuznetsova et al., 2017) in R (R Core Team, 2025). To further examine the plausibility effect within each secondary-task condition, follow-up pairwise comparisons of Implausible versus Plausible sentences were conducted separately for the NoSecondary and Articulation conditions, using estimated marginal means implemented in the emmeans package (Lenth and Piaskowski, 2026). To account for the two comparisons, Bonferroni correction was applied, and Bonferroni-adjusted *p* values were reported.

We included trials with correct and incorrect responses in this main analysis. We additionally conducted an analysis restricted to correct responses only. In summary, we created four models by manipulating onset time (the verb vs. the beginning of the sentence) and trial type (all trials vs. correct responses only). In the results section, we mainly report results from models with verb onset including all trials. Please see the Supplementary Material, Section 3 for all the results.^2^

### 2.7. Individual differences

Finally, we conducted an exploratory analysis of individual differences in WMC and their effects on both behavioral accuracy and brain activity. For behavioral data, within each condition, we tested whether WMC predicted correctness, using the GLM model (with random intercepts for participants and items).

For brain activity, we constructed first-level contrast images for each condition relative to the other conditions and entered these into group-level models, including WMC as a covariate. To assess both positive and negative relationships between WMC and brain activity, we tested separate contrasts for positive and negative effects of the covariate. This analysis identified voxels where brain activity during the task was significantly correlated with WMC across participants. To restrict our search to the LIFGtri and LIFGorb, we applied a priori mask (the same masks as used in the ROI analysis). We set a voxel-level FWE-corrected threshold of *p <* 0.05 as the significance level.

## 3. Results

### 3.1. Accuracy

Figure 2 represents the comprehension accuracy for each condition. The GLM model reveals that there were significant main effects of secondary task and plausibility: accuracy was lower in the Articulation condition than in the NoSecondary condition (*β* = *−*0.69, SE = 0.21, *z* = *−*3.30, *p <* 0.001), and lower in the Implausible condition than in the Plausible condition (*β* = *−*1.12, SE = 0.24, *z* = *−*4.62, *p <* 0.001). We did not find a significant interaction between the two factors (*β* = 0.46, SE = 0.26, *z* = 1.77, *p* = 0.078).

**Figure 2:**
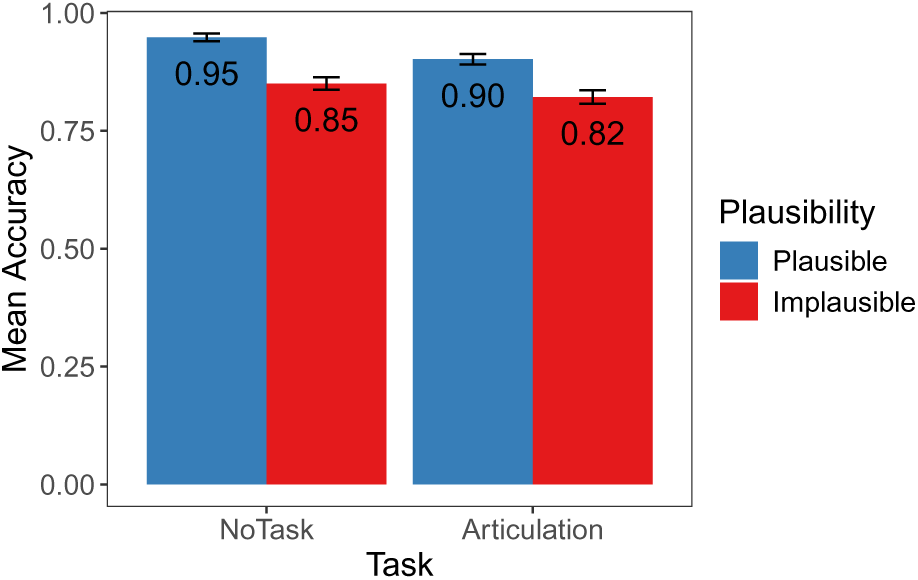
Accuracy of each condition. Accuracy was lower in the Articulation condition than in the NoSecondary condition, and lower in the Implausible condition than in the Plausible condition. There is no interaction. The error bar represents standard errors.

### 3.2. Imaging results

#### 3.2.1. Main analyses

The whole-brain analysis revealed significant main effects of the secondary task and plausibility, but no interaction between them (see Figure 3 and Table 2). Specifically, the Articulation condition elicited significantly greater activation than the NoSecondary condition across most of the brain, including the bilateral postcentral gyri and extending to the bilateral precentral gyri, bilateral IFG, superior frontal gyrus (SFG), superior temporal gyrus, and supramarginal gyrus. The widespread activation observed in the Articulation condition likely reflects motor and articulatory processes associated with the secondary task. The Implausible condition elicited significantly greater activation than the Plausible condition in the bilateral IFG pars orbitalis, bilateral insula, and the medial part of the SFG (mSFG).

**Figure 3:**
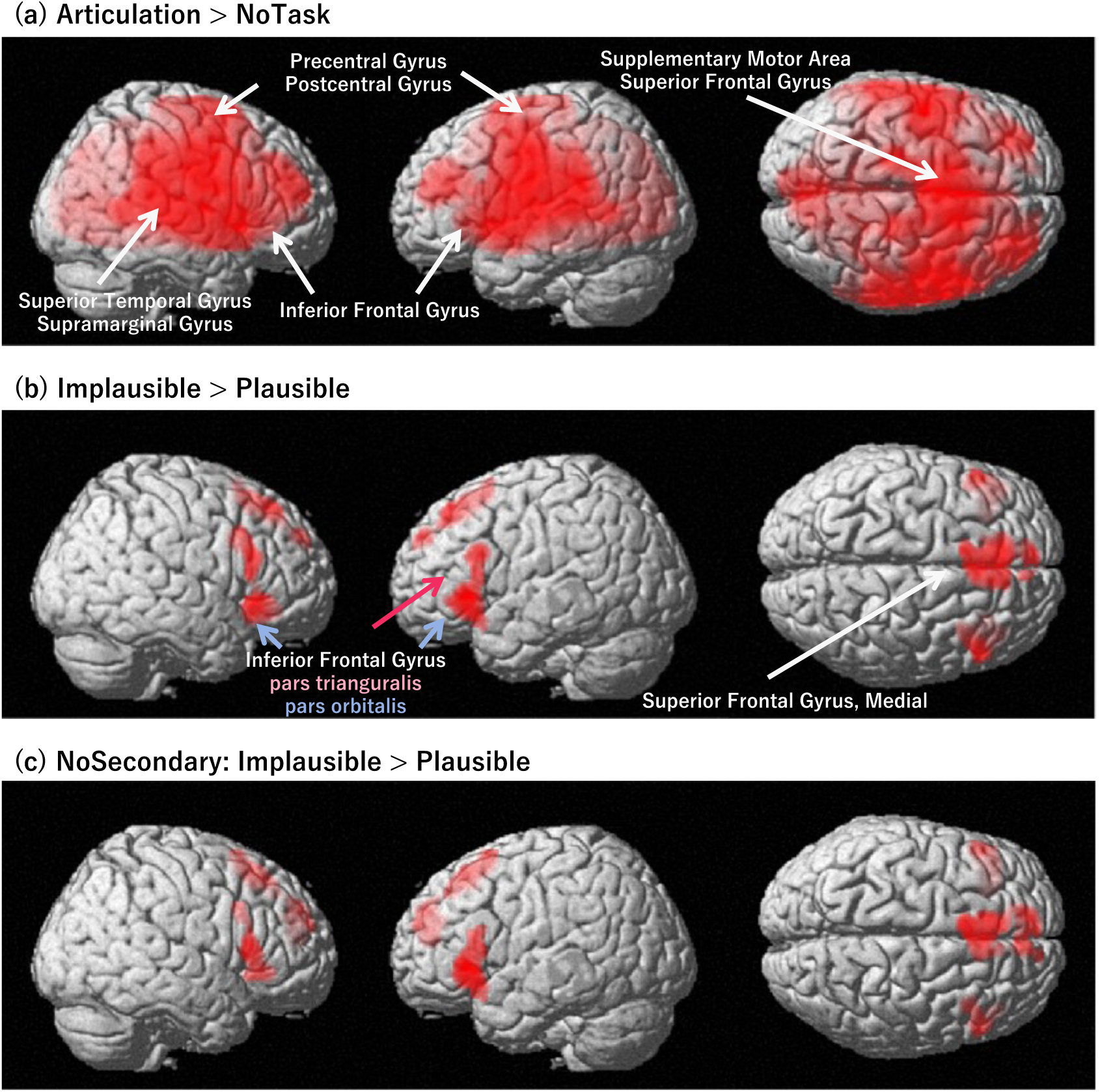
Results of the whole-brain analysis. Brain activity showing (a) greater activity in the Articulation condition compared to the NoSecondary condition, (b) greater activity in the Implausible condition than the Plausible condition, and (c) greater activity in the Implausible condition than the Plausible condition within the NoSecondary task.

**Table 2:**
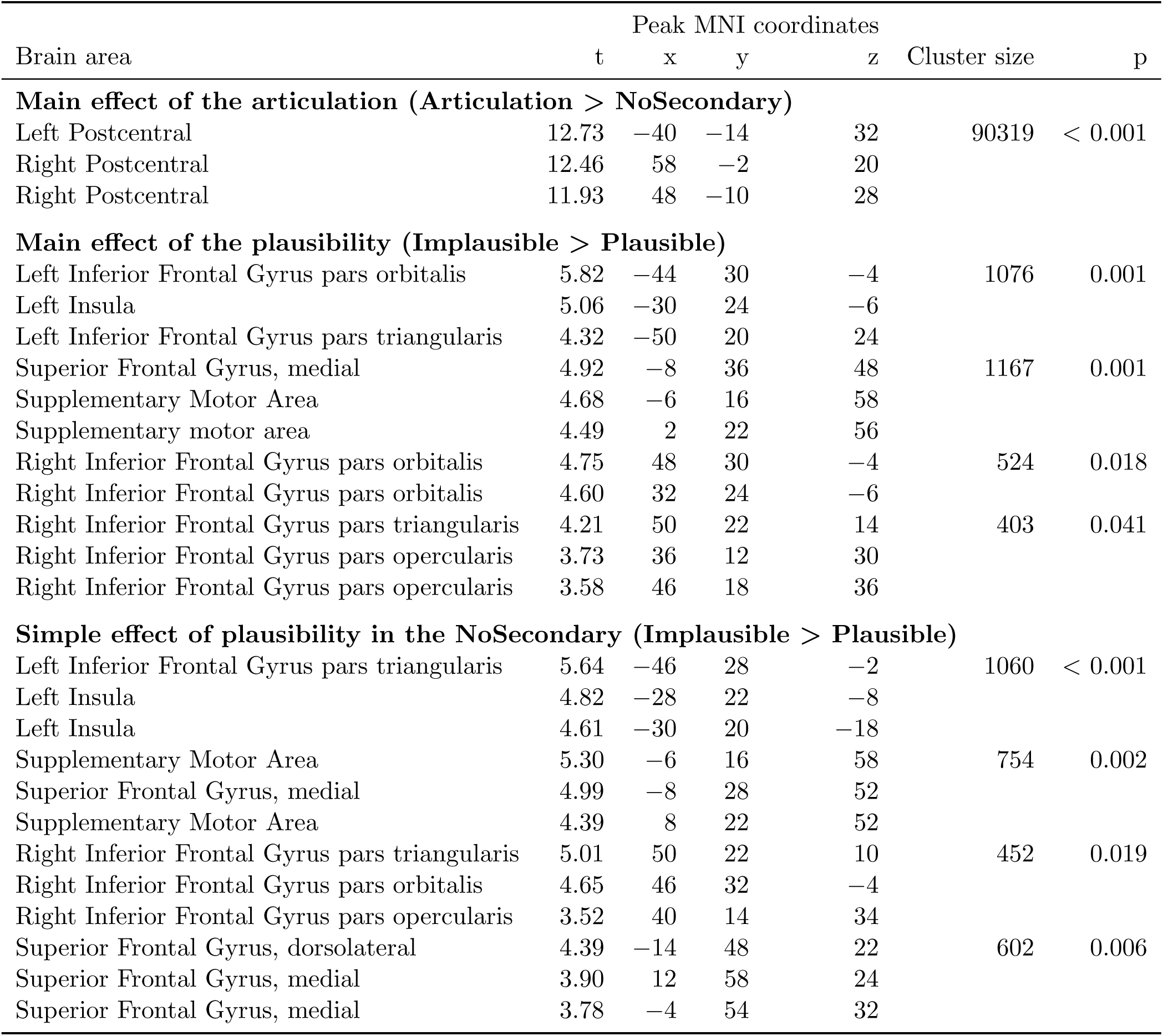
Significantly activated brain regions of whole brain analysis. For each area, the coordinates (x, y, z) of the activation peak in MNI space, peak t-value, and size of the activated cluster in number (k) of voxels are shown. FWE-corrected *p <* 0.05 with cluster-level after the initial threshold *p <* 0.001 with voxel-level.

Regarding the plausibility effect within each task, within the NoSecondary condition, the Implausible condition elicited greater activation than the Plausible condition in the same regions identified in the main effects (see Figure 3 and Table 2). In contrast, within the Articulation condition, no significant difference between the Plausible and Implausible conditions was found.

As shown in Section 2 of the Supplementary Material, other models also showed patterns similar to the main results. Specifically, when the analysis was limited to trials with correct responses, plausibility effects in the bilateral IFG and mSFG were seen. The two whole-sentence models also showed greater activation in the left IFG and mSFG.

In the ROI analysis, the LME models reveal asymmetric results between LIFGtri and LIFGorb (see Figure 4). In both regions, the Implausible elicited greater activation than the Plausible condition (LIFGtri: *β* = 0.68, SE = 0.25, *t* = 2.76, *p* = 0.006; LIFGorb: *β* = 0.74, SE = 0.24, *t* = 3.11, *p* = 0.002). However, LIFGtri showed greater activation in the Articulation condition than in the NoSecondary condition (*β* = 0.82, SE = 0.25, *t* = 3.33, *p* = 0.001), whereas no significant effect of the secondary task was observed in LIFGorb (*β* = 0.31, SE = 0.24, *t* = 1.32, *p* = 0.190). No significant interaction between plausibility and the secondary task was observed in either region (LIFGtri: *β* = *−*0.27, SE = 0.35, *t* = *−*0.77, *p* = 0.443; LIFGorb: *β* = *−*0.26, SE = 0.34, *t* = *−*0.77, *p* = 0.445).

**Figure 4:**
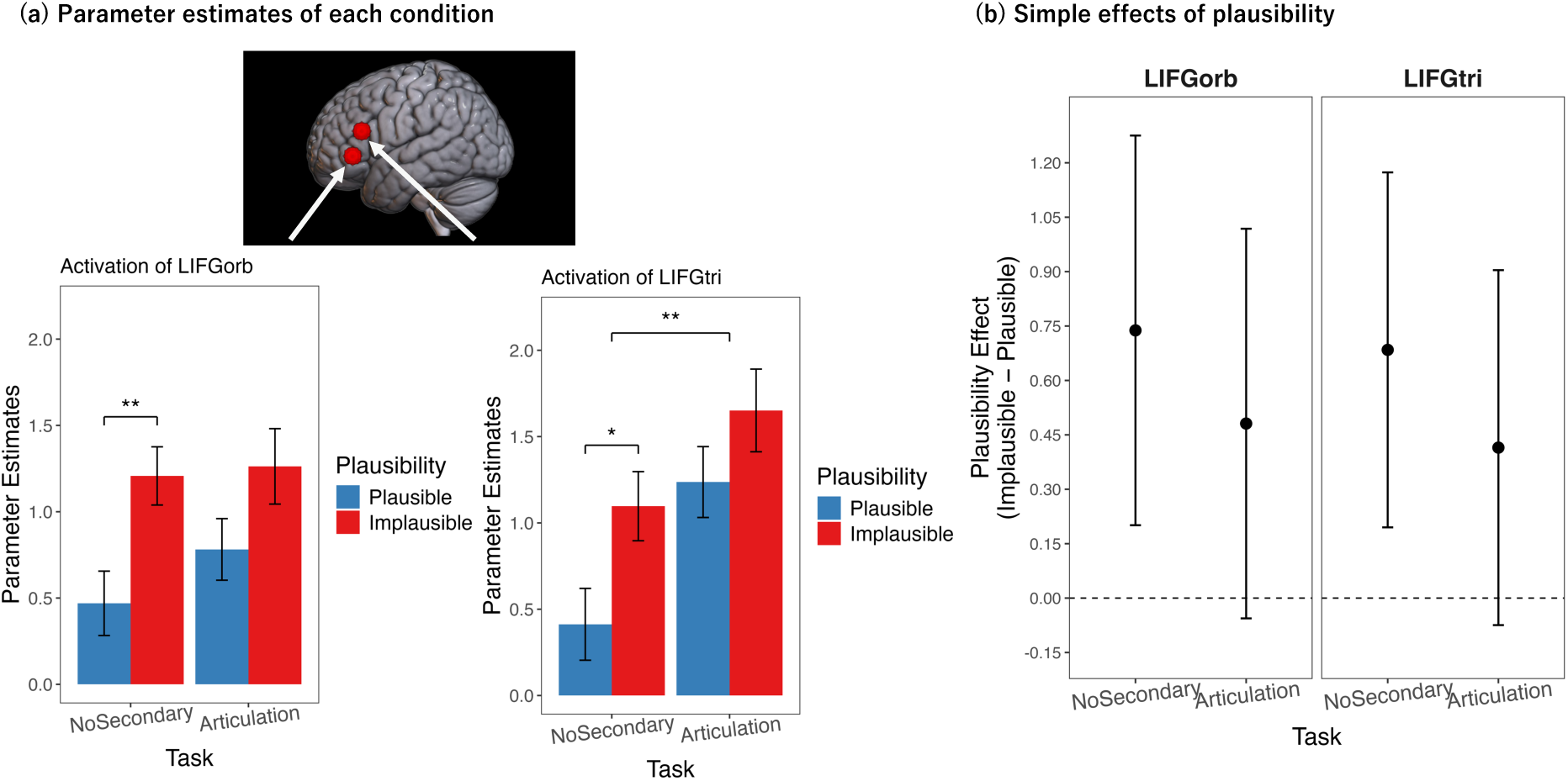
Results of the regions of interest (ROI) analyses. (a) ROI definition and parameter estimates in LIFGtri and LIFGorb. ROIs used in the analysis, including the left inferior frontal gyrus pars triangularis (LIFGtri) and pars orbitalis (LIFGorb), shown in red. The graph shows mean parameter estimates extracted from each ROI for each experimental condition. Error bars indicate standard errors. Both regions showed greater activation for Implausible than Plausible sentences. However, only LIFGtri exhibited increased activation in the Articulation condition relative to the NoSecondary condition, whereas no significant effect of the secondary task was observed in LIFGorb. No significant interaction between plausibility and secondary task was found in either region. (b) Simple effects of plausibility within each task. In both regions, the plausibility effects in the Articulation task were numerically lower than those in the NoSecondary task.

Post-hoc tests show the estimated marginal means between the Plausible and Implausible conditions in each task. Within the NoSecondary task, the Implausible condition elicited greater activation than the Plausible condition, both in LIFGtri (Estimate = 0.68, 95% Confidence Interval (95% CI) = [0.12, 1.25], *t* = 2.76, adjusted *p* = 0.013) and in LIFGorb (Estimate = 0.74, 95% CI = [0.20, 1.28], *t* = 3.11, adjusted *p* = 0.004). However, there is no significant plausibility effect within the Articulation task, either in LIFGtri (Estimate = 0.42, 95% CI = [*−*0.15, 0.98], *t* = 1.67, adjusted *p* = 0.192) or in

There is no significant Task *×* Plausibility interaction or simple plausibility effect within the Articulation task. FWE-corrected *p <* 0.05 with cluster-level after the initial threshold *p <* 0.001 with voxel-level in (a–b). FWE-corrected *p <* 0.025 with cluster-level after the initial threshold *p <* 0.001 with voxel-level in (c).

LIFGorb (Estimate = 0.48, 95% CI = [*−*0.06, 1.02], *t* = 2.03, adjusted *p* = 0.089).^3^ Thus, the simple effect of plausibility is numerically lower in the Articulation task than in the NoSecondary condition (see Figure 4b)

#### 3.2.2. Exploratory analyses

Although the main effect of plausibility was significant and the plausibility *×* task interaction was not significant, the plausibility effect was significant in the NoSecondary condition but not in the Articulation condition. We therefore conducted exploratory equivalence tests to assess whether the plausibility *×* task interaction and the plausibility effect under Articulation could be considered practically equivalent to zero. Using the two one-sided tests (TOST) procedure (Caldwell, 2026; Lakens, 2017), neither effect met the criterion for equivalence in either LIFG ROI. Thus, this study did not provide sufficient evidence to conclude that these effects were negligibly small. Details of these exploratory analyses are reported in the Section 4 of the Supplementary Material.

Finally, we conducted a simulation-based power analysis to assess the sample size required to detect the plausibility *×* task interaction under the variability observed in the present data. We considered an interaction of *−*0.23, approximately equal to the observed effect, and a larger interaction of *−*0.50. Under the reference heterogeneity scenario, the simulated power to detect an interaction of *−*0.23 at the present sample size (N = 58) was only 10.5% for the LIFGtri and 11.4% for the LIFGorb. Power exceeded 0.80 at approximately N = 800–1,600 for an interaction of *−*0.23 and at approximately N = 200–400 for an interaction of *−*0.50, depending on the assumed degree of between-participant heterogeneity. These results indicate that the present sample had limited sensitivity to an interaction of the magnitude observed here. Details of the simulation are provided in Section 5 of the Supplementary Material.

### 3.3. Individual differences

No significant WMC-related effects were observed in any condition for either positive or negative contrasts. Only in the model for the verb region that includes only correct trials, higher WMC scores were associated with reduced activation in the LIFGtri (voxel-level FWE-corrected *p* = 0.045, *t* = 3.06, MNI coordinates: [*−*46, 28, 16], cluster size: 2). See Section 3.5 of the Supplementary Material for details. Likewise, in the behavioral analysis, we did not find a significant effect of WMC scores on accuracy across conditions.

## 4. Discussion

The present study examined whether phonological-loop constraints modulate the use of plausibility information during thematic-role comprehension. Specifically, we tested whether articulatory suppression alters behavioral and neural responses to sentences in which syntactically assigned thematic roles are implausible given world knowledge. Three main findings emerged. First, we found no clear evidence that articulatory suppression modulated the plausibility effect. Second, irrespective of this uncertainty about task modulation, at least in the single-task condition, implausible sentences robustly elicited greater neural responses than plausible sentences, particularly in the LIFG and mSFG. Third, although LIFGtri and LIFGorb were both sensitive to thematic-role plausibility, only LIFGtri showed greater overall activation during concurrent articulation, suggesting the possibility that these subregions differ in their sensitivity to increased task demands. We discuss these findings in detail below.

### 4.1. Relationship between comprehension based on world knowledge and working memory constraints

The central question of this study is whether the phonological loop, one of the components of working memory, contributes to the use of plausibility information during thematic-role comprehension. As outlined in the Introduction, if the phonological loop helps maintain incoming linguistic information during the integration of world knowledge and input, articulatory suppression is expected to reduce the difference between plausible and implausible sentences.

However, the present results provide no clear evidence that the the articulatory-suppression task modulated the plausibility effect. Across tasks, implausible sentences elicited greater neural responses than plausible sentences in the LIFG and mSFG, whereas the interaction between plausibility and task was not significant in either the whole-brain analysis or the ROI analyses of the LIFG. Thus, the present data do not demonstrate that constraining the function of the phonological loop changes neural sensitivity to thematic-role plausibility.

Importantly, however, the absence of a significant interaction should not be interpreted as evidence that the plausibility effect was preserved to the same extent across the NoSecondary and Articulation conditions. In the follow-up analyses, significant Implausible *>* Plausible contrasts were observed in the NoSecondary condition, whereas corresponding effects in the Articulation condition did not survive correction for multiple comparisons in either the whole-brain or the ROI analyses. These findings therefore leave open the question of whether the plausibility effect is preserved under concurrent articulation.

As illustrated in Figure 4, the plausibility effect was numerically smaller in the Articulation condition than in the NoSecondary condition, which is consistent with the idea that phonological-loop constraints attenuated the plausibility effect. However, the interaction was not statistically significant, and the equivalence tests did not provide evidence that the difference between the two task conditions was negligible. Likewise, the analyses did not establish that the plausibility effect during articulation was equivalent to zero.

The present results, therefore, do not provide sufficient evidence that either thematic-role plausibility processing was remained unchanged during concurrent articulation or it was eliminated by the secondary task. One likely reason for this uncertainty is the substantial variability in the estimated task-by-plausibility interaction. Exploratory power calculations indicated that increasing the sample size and reducing between-participant variability would allow future studies to obtain more precise estimates of the interaction. However, given an effect of the magnitude observed in the present study, the required sample size would remain very large; even under assumptions of relatively low variability, approximately 800 participants would be required. The present experiment may therefore have been insufficiently sensitive to determine whether concurrent articulation produces a small attenuation of the plausibility effect.

The absence of a clear interaction is unlikely to reflect a complete failure of the secondary task to impose additional processing demands. Behavioral accuracy decreased under the dual-task condition. In addition, concurrent articulation elicited increased activation in the supramarginal gyrus, a region associated with auditory and somatosensory feedback control, as well as in the IFG and mSFG, which have been implicated in verbal working memory and sentence processing (Kearney and Guenther, 2019). Thus, these findings suggest that the articulation task placed additional demands on working-memory functions.

The present findings can also be considered in relation to previous work on syntactic processing using a similar dual-task approach. Rogalsky et al. (2008) reported that articulatory suppression reduced the syntactic-complexity effect in LIFGoper, whereas the effect remained in LIFGtri. By contrast, this study did not observe a reduced plausibility effect under articulatory suppression in either the LIFGtri or the LIFGorb. This difference does not demonstrate that the use of world knowledge is independent of the phonological loop. Rather, it indicates that the task-dependent modulation observed for syntactic complexity in previous work was not clearly demonstrated for thematic-role plausibility under the present experimental conditions.

To summarize, the present findings do not provide conclusive evidence that the phonological loop supports the processing of plausibility information based on world knowledge. However, the current data are not sufficiently precise to rule out a smaller attenuation of plausibility effect. Therefore, the role of phonological loop in thematic-role interpretation with world knowledge remains unresolved rather than demonstrably absent.

### 4.2. Robustness of the plausibility effect in LIFG and mSFG

Although the effect of concurrent articulation on thematic-role plausibility remains uncertain, the present study showed reliable neural sensitivity to thematic-role plausibility. A main effect of plausibility was observed across tasks, with robust Implausible *>* Plausible effects also found within the NoSecondary condition. Implausible sentences were associated with lower comprehension accuracy and elicited greater activation than plausible sentences in regions including the LIFGorb, LIFGtri, and mSFG (including the supplementary motor area). These findings are consistent with previous studies reporting neural sensitivity to inconsistencies between syntactically specified interpretations and plausibility derived from world knowledge in Japanese, English, Mandarin Chinese, and Dutch (Hagoort et al., 2004; Kuperberg et al., 2008; Sassa et al., 2007; Tyler et al., 2010; van de Meerendonk et al., 2013; Ye and Zhou, 2009). We also observed greater activation in the insula, which has been argued to play an integrative role in sentence production, sentence comprehension, and emotion (Ardila et al., 2014; Gasquoine, 2014; Oh et al., 2014).

These findings can be interpreted in relation to the role of world knowledge in thematic-role comprehension. Comprehending implausible sentences imposes additional processing demands relative to plausible sentences. The present data provide further evidence that the neural network involving the insula, LIFG, and mSFG contributes to the processing of plausibility information used in thematic-role comprehension (Ardila et al., 2014).

Importantly, the involvement of the LIFG and mSFG was robust across several analytic choices. Greater responses to implausible than to plausible sentences in these regions were observed when the analysis window was restricted to the critical verb as well as when it encompassed the entire sentence. A similar spatial pattern was also obtained when analyses included all trials and when they were restricted to trials with correct behavioral responses. Thus, the observed LIFG and mSFG effects are unlikely to depend on a particular temporal window or on the inclusion of trials with incorrect responses. Instead, the convergence across these analyses provides robust evidence that these regions are sensitive to the plausibility derived from world knowledge.

### 4.3. Functional asymmetry within the LIFG

The LIFG has been a controversial region in terms of the relationship between working memory and sentence processing (Fedorenko et al., 2012; Fedorenko and Blank, 2020; Matchin et al., 2024; Rogalsky and Hickok, 2011). Although our data show the independent effect of sentence plausibility not modulated by the articulation task, ROI analyses suggest a possible functional asymmetry within the LIFG according to overall working memory demands. Our data show both LIFGtri and LIFGorb were sensitive to thematic-role plausibility. However, these regions did not activate uniformly in response to the task demands; only LIFGtri, not LIFGorb, showed significantly greater activity in the articulation task than in the single task.

This pattern suggests that the two regions may differ in their sensitivity to increased task demands. To test this possibility, we conducted an exploratory post-hoc analysis in which we combined data from the two ROIs and fitted the LME model with ROI and task as factors.^4^

One possible interpretation is that LIFGtri is more sensitive to modulation of brain activity to overall cognitive resource limitations, whereas LIFGorb may play a more stable role in sentence-level interpretation. The relatively weaker task effect in LIFGorb is consistent with previous work showing that this region contributes to the distinction between sentences and non-words, but not necessarily to other higher cognitive functions, including verbal working memory, spatial working memory, mathematics, and Stroop tasks (Fedorenko et al., 2011). In contrast, Fedorenko et al. (2011) reported that the superior-posterior part of the LIFG overlaps between sentence-sensitive regions and verbal working-memory regions. This finding is compatible with the present observation that LIFGtri was sensitive to overall task demands. Meta-analytic evidence is also broadly consistent with this view (Bulut, 2023).

A caution is needed because the critical interaction between plausibility and the secondary task was not significant. This interpretation should be taken as evidence for differential task sensitivity, rather than direct evidence that task demands changed plausibility processing.

To summarize, the present ROI findings suggest that LIFGtri and LIFGorb both contribute to the usage of world knowledge in thematic-role-interpretation, but only LIFGtri showed an exploratory association with increased task demands. This asymmetry is compatible with the possibility that subregions within the LIFG may play partially distinct roles: LIFGorb may support relatively stable sentence-level interpretation, whereas LIFGtri may be more sensitive to overall cognitive resource limitations during sentence comprehension.

### 4.4. Limitations and future directions

First, caution is warranted regarding functional localization within the LIFG because the functional role of LIFG remains controversial (Fedorenko et al., 2012; Fedorenko and Blank, 2020; Matchin et al., 2024; Rogalsky and Hickok, 2011). For example, some studies have argued that LIFGtri is selectively involved in syntactic processing (Rogalsky et al., 2008; Rogalsky and Hickok, 2011; Matchin, 2018). Other work has suggested that the localization of language-selective regions varies across individuals (Fedorenko et al., 2012; Fedorenko and Blank, 2020). This variability highlights the importance of considering individual differences and using participant-specific functional localization when evaluating how working memory resources shape neural responses during sentence comprehension.

Second, the interpretation of the dual-task manipulation remains limited. Concurrent articulation has been commonly used to constrain the phonological loop for a long time (e.g., Baddeley, 1979; Beni et al., 2005; Besner et al., 1981; Hitch and Baddeley, 1976; Slowiaczek and Clifton, 1980), but it might not be a pure phonological-loop manipulation. As shown by the broad activation in motor and somatosensory regions, concurrent articulation also engages speech-motor control, auditory and somatosensory feedback, and domain-general task coordination. Therefore, the present manipulation cannot completely isolate the phonological loop from other articulatory and executive demands. Future work using alternative load manipulations, such as nonverbal secondary tasks, articulatory suppression with different speech materials, or arithmetic problem solving, may help clarify which components of working memory contribute to sentence processing.

### 4.5. Conclusion

In conclusion, the present study links two types of accounts of sentence comprehension: accounts of how linguistic information and prior world knowledge jointly contribute to interpretation, and accounts of the working-memory architecture that supports online sentence processing. To examine this relationship, we used an fMRI dual-task paradigm in which resources of the phonological loop, a component of working memory, were constrained by concurrent articulation while participants comprehended sentences differing in thematic-role plausibility. Three main findings emerged. First, we found no clear evidence that concurrent articulation modulated the thematic-role plausibility effect. Second, we observed robust behavioral and neural sensitivity to plausibility, with implausible sentences being associated with lower comprehension accuracy and greater neural responses particularly in the LIFG and mSFG. Third, although both LIFGtri and LIFGorb were sensitive to plausibility, only LIFGtri showed increased activation under concurrent articulation, suggesting the possibility that these subregions may differ in their sensitivity to overall task demands. Together, these results help clarify how working-memory constraints shape the neural basis of sentence comprehension, particularly the use of plausibility information derived from prior world knowledge during the interpretation of who did what to whom.

## Author Contributions

RE: Conceptualization, Formal analysis, Funding acquisition, Investigation, Methodology, Project administration, Software, Visualization, Writing – original draft; HJ: Funding acquisition, Investigation, Methodology, Resources, Software, Supervision, Validation, Writing – review & editing; YK: Conceptualization, Supervision, Writing – review & editing; RK: Formal analysis, Investigation, Writing – review & editing; KM: Funding acquisition, Resources, Supervision, Writing – review & editing.

## Acknowledgments

I would like to thank Liwei Tan and Wang Yucheng (Tohoku University) for supporting data collection and Taiga Naoe (Showa Medical University, Tohoku University) for valuable feedback. This work was supported by JSPS KAKENHI Grant Numbers JP23KJ0199 and Fund for fMRI Research Challenge and Collaboration (CogNAC, Tohoku University).

## Footnotes

1 Response times were not part of the original analysis plan and were not recorded due to a technical error. Therefore, response-time data were unavailable for analysis.

2 In addition to the brain activities during sentence comprehension, we analyzed those during comprehension questions in accordance with a reviewer’s suggestion (See the Supplementary Material, Section 7).

3 Only in the model with the whole-sentence time window including only correct responses reveals the significance difference in plausibility within the Articulation task in LIFGtri. See the Supplementary Material, Section 3.5.

4 The model was specified as *Activation ∼ ROI ∗Task* +*Plausibility* +(1*|Participant*). Full results of this exploratory analysis are reported in the Supplementary Materials, Section 6. The analysis revealed a significant interaction between ROI and task (*β* = 0.51, SE = 0.25, *t* = 2.06, *p* = 0.041). A simple-effects analysis further showed a significant increase in activation during the Articulation condition relative to the NoSecondary condition in LIFGtri (Estimate = 0.69, 95% CI = [0.35, 1.03], *t* = 3.96, Bonferroni adjusted *p <* 0.001), but not in LIFGorb (Estimate = 0.18, 95% CI = [*−*0.16, 0.53], *t* = 1.06, Bonferroni adjusted *p* = 0.291).

